# Antifungal susceptibility profile of clinically relevant species of the genus *Sporothrix:* establishment of Epidemiological Cutoff Values (ECVs) according to CLSI broth microdilution methodology

**DOI:** 10.64898/2025.12.19.695392

**Authors:** Amanda R. dos Santos, Alexandro Bonifaz, Amanda Bombassaro, Ana Caroline de Sá Machado, Anderson M. Rodrigues, Andrew M. Borman, Arunaloke Chakrabarti, Anuradha Chowdary, Bruno Rediguieri, Elizabeth M. Johnson, Ferry Hagen, Flavio Queiroz-Telles, Gloria M. Gonzalez, Guillermo Garcia-Effron, Isabella Dib Ferreira Gremião, Jacques F. Meis, Juliana Possato Fernandes Takahashi, Luana Pereira Borba dos Santos, Marcia S. C. Melhem, Paola Cappellano, Raimunda Sâmia Nogueira Brilhante, Sandro Antonio Pereira, Sarah Santos Gonçalvez, Sarah Kidd, Sean X. Zhang, Shivaprakash Rudramurthy, Sonia Rozental, Susana Cordoba, Theun de Groot, Wei Liu, Nathan P. Wiederhold, Eelco F.J. Meijer, Shawn R. Lockhart, Philippe J. Dufresne

## Abstract

Sporotrichosis is a globally distributed subcutaneous mycosis caused mainly by *Sporothrix brasiliensis*, *S. schenckii*, and *S. globosa*. Cat-transmitted sporotrichosis, primarily caused by *S. brasiliensis* in South America and to a lesser extent by *S. schenckii* in Southeast Asia, is emerging as a significant public health concern, due its outbreak potential. Itraconazole is the first-choice drug for treatment of human and cats, but reduced susceptibility has been reported based on previously proposed epidemiological cut-off values (ECVs). To support resistance surveillance, we aimed to establish CLSI-endorsed ECVs for these clinically relevant *Sporothrix* species. A total of 3,588 minimum inhibitory concentration (MIC) values for seven antifungal agents (amphotericin B, itraconazole, posaconazole, voriconazole, isavuconazole, olorofim, and terbinafine) were obtained from 19 international laboratories. Four of seven antifungals met the CLSI M57 guidelines criteria to determine the ECV. Established ECVs for amphotericin B were found to be high with 8 µg/mL for *S. brasiliensis* and *S. globosa*, and 4 µg/mL for *S. schenckii*. Itraconazole ECVs were 4 µg/mL for *S. brasiliensis* and *S. schenckii*. Posaconazole ECVs were 4 µg/mL for all three species (tentative for *S. globosa*), while the terbinafine ECV for *S. brasiliensis* was 0.12 µg/mL. Olorofim demonstrated good *in vitro* activity, particularly against *S. brasiliensis*. Overall, this study establishes validated ECVs for key antifungals against *Sporothrix* species and identifies a low prevalence of non-wild type (NWT) isolates, supporting ongoing antifungal resistance monitoring.

## Introduction

Sporotrichosis is a ubiquitous cutaneous/subcutaneous mycosis caused by pathogenic species of the *Sporothrix* genus (1). It is more prevalent in tropical and subtropical areas, where infections mostly occur by percutaneous inoculation from plant material containing *Sporothrix* spp., the classical form of the disease (2). Outside South America this is the common form of the disease and is mainly caused by *S. schenckii* and *S. globosa* (3). In the last three decades a new species in the *Sporothrix* pathogenic clade*, S. brasiliensis*, has emerged in Brazil and thereafter in other countries in South America (4). *S. brasiliensis* causes large outbreaks in the cat population and is increasingly transmitted to humans by direct contact with infected cats (4). In humans, the usual clinical presentations of sporotrichosis are fixed cutaneous or lymphocutaneous, and multifocal disseminated infections in HIV patients. Other clinical presentations including osteoarticular, pulmonary, and multifocal infections in non-immunocompromised patients, are relatively rare (5, 6). In cats, the most prevalent clinical presentation is the disseminated cutaneous form, typically associated with extracutaneous manifestations, including respiratory involvement and lymphadenomegaly (7,8). Remarkably, in the last decade there have also been several reports of cat-transmitted sporotrichosis in Thailand and Malaysia, which were caused by *S. schenckii* (9–11).

In South America, and to a lesser extent in Southeast Asia, zoonotic sporotrichosis has become a major public health problem, because of its capacity to rapidly spread and its epidemic potential (12). Currently, there is no vaccine against sporotrichosis and the treatment of infected cats is the main prevention strategy to prevent outbreaks and further spread of the disease (13). There are limited antifungal treatment options in cats and humans. Itraconazole is the first-choice antifungal drug to the human disease, while terbinafine and potassium iodide are used as alternatives (14). Amphotericin B is used for severe human infections (15, 16). In feline sporotrichosis, itraconazole administered either as monotherapy or in combination with potassium iodide, remains the first-line drugs. Deoxycholate amphotericin B can be administered in the lesion in cases refractory to itraconazole (17). Terbinafine demonstrates limited antifungal effectiveness (18). Additionally, there are few documented cases reporting successful use of posaconazole and isavuconazole in treatment-refractory infections (19,20). In Brazil, effective treatment of feline sporotrichosis is hindered by multiple factors, including owner treatment non-adherence, social-economic constraints, the long duration of therapy and the cost of itraconazole, which is not provided free of charge in many public health services. (21). Furthermore, challenges with the administration of the antifungal in cats have been documented, and treatment abandonment has been reported in 34-39% of infected cats (22, 23). Poor treatment compliance and treatment failure in cats may have an impact on human health, as it can lead to increase fungal burden, antifungal exposure and could favor the development of antimicrobial resistance (AMR). Reduced susceptibility of *S. brasiliensis* to itraconazole was suggested in a human case non-responsive to itraconazole (24). As cats are treated with the same antifungal agents used in humans, the potential emergence of antimicrobial resistance may represent an additional challenge for the management of human sporotrichosis.

The detection of isolates with non-wild type (NWT) minimal inhibitory concentrations (MICs) enables surveillance of the emergence and spread of antifungal resistance (25). Establishing a baseline of MICs and associated clinical presentations are the first steps for monitoring the emergence of resistance. For this purpose, antifungal breakpoints for medically relevant *Sporothrix* species are necessary. Owing to a lack of studies reporting MICs along with clinical data and treatment outcomes, breakpoints have not been established, nor are official epidemiological cut-off values (ECVs) available, which enable the identification of isolates with reduced susceptibility and potential antifungal resistance mechanisms.

Several studies have reported azole MIC values for *S. brasiliensis* (26–31), whereas a multicenter international study proposed ECVs for *S. brasiliensis* and *S. schenckii* to azoles and amphotericin B, and to terbinafine for *S. brasiliensis* only (27). Although this study included a high number of MIC values, results were not submitted for approval and validation to the Clinical and Laboratory Standards Institute (CLSI). Here, using an expanded contemporary dataset from a total of 19 participating laboratories, including some of the MICs values presented in the previous ECV study (27), we established MIC distributions, ECVs, percent non-wild-type (NWT), modal MICs, MIC_90_s, and geometric mean MICs for seven antifungals including amphotericin B, olorofim, terbinafine and triazoles (itraconazole, posaconazole, voriconazole and isavuconazole) for *S. schenckii, S. globosa* and/or *S. brasiliensis*.

## Results

A total of 3,588 MIC data for *S. brasiliensis*, *S. globosa*, and *S. schenckii* isolates were obtained from 19 laboratories who performed AFST using the broth microdilution method as outlined in the CLSI reference standard M38 Ed3 for filamentous fungi (32). A subset of the MICs (up to 2017) came from nine laboratories involved in a previous study (27). Additional contemporary MIC datasets were also received from some of those laboratories as well as from ten new sites. MIC distributions were determined for amphotericin B (**Table 1**), olorofim, terbinafine (**Table 2**), and triazoles (**Table 3**), together with the geometric mean MIC, MIC_50_ and MIC_90_ data. The number of isolates and participating laboratories varied across antifungals, as did the number of isolates among the three species (**Table 4**). MIC datasets were rejected if they did not conform to CLSI M57 guidelines (such as being multimodal, truncated at the highest or lowest value, or not within 2 dilutions of the central modal MIC of most laboratories). More significant interlaboratory variation and rejected MIC datasets from laboratories were seen with *S. brasiliensis* for amphotericin B (22%) and azoles (17-30%), and *S. schenckii* with amphotericin B (31%) **(Table 4)**. For all other species/antifungal combinations, the percentage of MIC datasets from laboratories included was ≥85%.

**Table 1.**
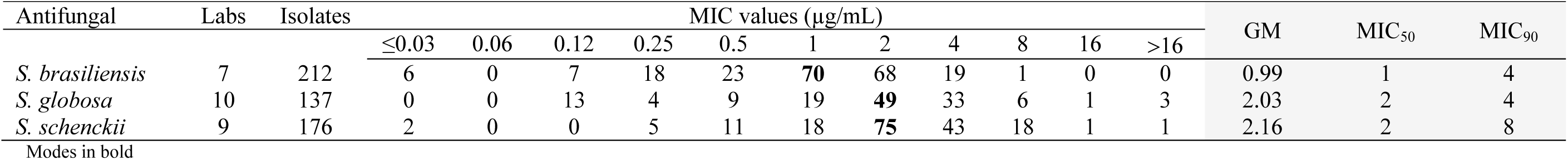
Amphotericin B MIC distribution for *Sporothrix* species.

**Table 2.** Olorofim and terbinafine MIC distributions for *Sporothrix* species.

| Antifungal | Labs | Isolates | MIC values (µg/mL) |  |  |  |  |  |  |  |  |  |  |  | GM | MIC<sub>50</sub> | MIC<sub>90</sub> |
| --- | --- | --- | --- | --- | --- | --- | --- | --- | --- | --- | --- | --- | --- | --- | --- | --- | --- |
|  |  |  | 0.004 | 0.008 | 0.016 | 0.03 | 0.06 | 0.12 | 0.25 | 0.5 | 1 | 2 | 4 | 8 |  |  |  |
| **Olorofim** |  |  |  |  |  |  |  |  |  |  |  |  |  |  |  |  |  |
| *S. brasiliensis* | 1 | 52 | 4 | 9 | **14** | 13 | 3 | 8 | 2 | 0 | 0 | 0 | 0 | 0 | 0.24 | 0.016 | 0.12 |
| *S. globosa* | 1 | 33 | 0 | 0 | 0 | 0 | 0 | 0 | 2 | 14 | **17** | 0 | 0 | 0 | 0.69 | 1 | 1 |
| *S. schenckii* | - | - | - | - | - | - | - | - | - | - | - | - | - | - | - | - | - |
| **Terbinafine** |  |  |  |  |  |  |  |  |  |  |  |  |  |  |  |  |  |
| *S. brasiliensis* | 4 | 270 | 1 | 5 | 9 | 87 | **129** | 27 | 6 | 0 | 0 | 0 | 0 | 0 | 0.05 | 0.06 | 0.12 |
| *S. globosa* | 3 | 38 | 0 | 0 | 0 | 0 | 4 | 0 | 2 | 15 | **17** | 0 | 0 | 0 | 0.53 | 0.5 | 1 |
| *S. schenckii* | 3 | 42 | 0 | 0 | 0 | 1 | 11 | **15** | 8 | 7 | 0 | 0 | 0 | 0 | 0.14 | 0.12 | 0.5 |
| Modes in bold |  |  |  |  |  |  |  |  |  |  |  |  |  |  |  |  |  |

Key: GM, geometric mean; MIC, minimum inhibitory concentration; MIC<sub>50</sub> or MIC<sub>90</sub>, MIC that inhibits 50% or 90% of the isolates tested.

**Table 3.** Triazole MIC distributions for *Sporothrix* species.

| Antifungal | Labs | Isolates | MIC values (µg/mL) |  |  |  |  |  |  |  |  |  |  |  | GM | MIC<sub>50</sub> | MIC<sub>90</sub> |
| --- | --- | --- | --- | --- | --- | --- | --- | --- | --- | --- | --- | --- | --- | --- | --- | --- | --- |
|  |  |  | ≤0.016 | 0.03 | 0.06 | 0.12 | 0.25 | 0.5 | 1 | 2 | 4 | 8 | 16 | >16 |  |  |  |
| **Isavuconazole** |  |  |  |  |  |  |  |  |  |  |  |  |  |  |  |  |  |
| *S. brasiliensis* | 3 | 93 | 0 | 0 | 1 | 1 | 2 | 1 | 9 | 14 | 20 | **35** | 9 | 1 | 4.06 | 4 | >16 |
| *S. globosa* | 3 | 20 | 0 | 0 | 0 | 0 | 0 | 0 | 0 | 1 | 2 | 4 | **11** | 2 | 11.71 | 16 | >16 |
| *S. schenckii* | 5 | 74 | 0 | 0 | 0 | 0 | 0 | 0 | 10 | 5 | 18 | **26** | 8 | 7 | 5.71 | 8 | >16 |
| **Itraconazole** |  |  |  |  |  |  |  |  |  |  |  |  |  |  |  |  |  |
| *S. brasiliensis* | 7 | 216 | 5 | 1 | 3 | 14 | 35 | 50 | **69** | 29 | 2 | 0 | 0 | 8 | 0.65 | 1 | 2 |
| *S. globosa* | 10 | 96 | 1 | 13 | 4 | 9 | 15 | **16** | 9 | 12 | 3 | 4 | 1 | 9 | 0.55 | 0.5 | >16 |
| *S. schenckii* | 11 | 242 | 0 | 2 | 6 | 10 | 31 | **67** | 63 | 38 | 5 | 2 | 1 | 17 | 0.86 | 1 | 4 |
| **Posaconazole** |  |  |  |  |  |  |  |  |  |  |  |  |  |  |  |  |  |
| *S. brasiliensis* | 6 | 291 | 0 | 0 | 2 | 4 | 13 | 58 | **100** | **100** | 11 | 0 | 0 | 3 | 1.08 | 1 | 2 |
| *S. globosa* | 8 | 80 | 0 | 1 | 0 | 3 | 14 | 19 | **27** | 8 | 2 | 0 | 1 | 5 | 0.69 | 1 | 4 |
| *S. schenckii* | 10 | 266 | 0 | 0 | 1 | 10 | 14 | 54 | **83** | 41 | 11 | 9 | 8 | 35 | 1.57 | 1 | >16 |
| **Voriconazole** |  |  |  |  |  |  |  |  |  |  |  |  |  |  |  |  |  |
| *S. brasiliensis* | 5 | 137 | 0 | 0 | 0 | 0 | 0 | 2 | 6 | 6 | 16 | 41 | **54** | 12 | 9.03 | 8 | 16 |
| *S. globosa* | 9 | 111 | 0 | 11 | 0 | 6 | 2 | 3 | 3 | 4 | 5 | 15 | 22 | **40** | 9.29 | 16 | >16 |
| *S. schenckii* | 10 | 266 | 0 | 4 | 0 | 0 | 1 | 4 | 13 | 27 | 43 | 65 | **86** | 23 | 6.83 | 8 | 16 |

Modes in bold Key: GM, geometric mean; MIC, minimum inhibitory concentration; MIC<sub>50</sub> or MIC<sub>90</sub>, MIC that inhibits 50% or 90% of the isolates tested.

**Table 4.** Epidemiological cutoff values (ECV) for *Sporothrix* species and % non-wildtype (NWT)

| Species | Labs<br>included out<br>of total | Isolates<br>included out<br>of total | ECV | %NWT |
| --- | --- | --- | --- | --- |
| **Amphotericin B** |  |  |  |  |
| *S. brasiliensis* | 7/9 (78%) | 212/370 (57%) | 8<sup>b</sup> | 0% |
| *S. globosa* | 10/10 (100%) | 137 (100%) | 8<sup>b</sup> | 3.1% |
| *S. schenckii* | 9/13 (69%) | 176/280 (63%) | 4<sup>b</sup> | 2.3% |
| **Itraconazole** |  |  |  |  |
| *S. brasiliensis* | 7/10 (70%) | 216/428 (51%) | 4<sup>b</sup> | 3.7% |
| *S. globosa* | 10/11 (91%) | 96/129 (74%) | ILV | - |
| *S. schenckii* | 11/13 (85%) | 242/286 (85%) | 4<sup>b</sup> | 8.3% |
| **Isavuconazole** |  |  |  |  |
| *S. brasiliensis* | 3/4 (75%) | 93/154 (60%) | TR-H | - |
| *S. globosa* | 3/3 (100%) | 19 (100%) | TR-H | - |
| *S. schenckii* | 5/5 (100%) | 74 (100%) | TR-H | - |
| **Posaconazole** |  |  |  |  |
| *S. brasiliensis* | 7/9 (78%) | 291/331 (88%) | 4<sup>b</sup> | 1.0% |
| *S. globosa* | 8/9 (89%) | 80/113 (71%) | (4)<sup>b</sup> | 7.5% |
| *S. schenckii* | 10/10 (100%) | 266 (100%) | 4<sup>b</sup> | 19.5% |
| **Voriconazole** |  |  |  |  |
| *S. brasiliensis* | 5/6 (83%) | 137/189 (100%) | TR-H | - |
| *S. globosa* | 9/9 (100%) | 111 (100%) | TR-H | - |
| *S. schenckii* | 10/10 (100%) | 266 (100%) | TR-H | - |
| **Olorofim** |  |  |  |  |
| *S. brasiliensis* | 1/1 (100%) | 52 (100%) | - | - |
| *S. globosa* | 1/1 (100%) | 33 (100%) | - | - |
| *S. schenckii* | - | - | - | - |
| **Terbinafine** |  |  |  |  |
| *S. brasiliensis* | 4/4 (100%) | 270 (100%) | 0.12 | 4.4% |
| *S. globosa* | 3/3 (100%) | 38 (100%) | (2)<sup>a</sup> | 0% |
| *S. schenckii* | 3/3 (100%) | 42 (100%) | (0.5)<sup>a</sup> | 0% |

Key: -, ECV not defined; TR-H, ECV cannot be defined for this species-antifungal combination because the MIC distribution is truncated at the high end of the recommended testing range. ILV, ECV cannot be defined for this species-antifungal combination because of interlab variation. <sup>a</sup>Tentative ECVs are presented in parentheses (the number of isolates is < 100 and >35). <sup>b</sup>The ECV is high which indicates limited *in vitro* susceptibility to this agent. An MIC lower than the ECV does not imply that the isolate is susceptible to that antifungal agent.

Data for four of seven antifungals met the CLSI M57 guidelines (minimum of 3 different laboratories and 100 isolates) for determination of ECVs via the iterative statistical method with ECOFFinder (V2.1). Tentative ECVs were also calculated if numbers of isolates were insufficient (<100 and >35 isolates). For amphotericin B, ECVs were 8 µg/mL for *S. brasiliensis* and *S. globosa*, and 4 µg/mL for *S. schenckii*. ECVs for itraconazole and posaconazole were 4 µg/mL for *S. brasiliensis* and *S. schenckii*, while a tentative ECV of 4 µg/mL calculated for *S. globosa* and posaconazole. Finally, the ECVs for terbinafine was 0.12 µg/mL for *S. brasiliensis*, and only tentative ECVs for *S. globosa* (2 µg/mL) and *S. schenckii* (0.5 µg/mL) could be defined due to lack of isolates, (**Table 4**). The itraconazole ECV for *S. globosa* could not be determined due to a high variability in MICs between different laboratories. Additionally, the ECVs for isavuconazole and voriconazole could not be established, as MICs were truncated on the high end of the recommended testing range. Last, olorofim susceptibility was only determined in one laboratory, where it showed good antifungal activity, especially toward *S. brasiliensis* compared with *S. globosa* (**Table 4**).

The percentage of NWT isolates with MICs above the established ECV was determined for each drug and species. For amphotericin B and terbinafine, the percentage of NWT strains was <5% for all species. The percentage of NWT strains was <5% for itraconazole and posaconazole against *S. brasiliensis* isolates. In contrast, 8.3% and 19.5% of *S. schenckii* isolates were NWT for itraconazole and posaconazole, respectively, whereas 7.5% of *S. globosa* isolates were NWT for posaconazole (**Table 4**).

## Discussion

In this study, we established ECVs for the filamentous phase of *S. brasiliensis*, *S. schenckii* and *S. globosa* for the antifungal drugs amphotericin B, itraconazole, terbinafine, and posaconazole following CLSI M57 guidelines. These ECVs were generated with a contemporary MIC dataset and accepted by the CLSI antifungal subcommittee. For terbinafine and posaconazole, tentative ECVs were proposed for *S. globosa* and/or *S. schenckii*. In addition, MIC data was analyzed for olorofim, which showed good antifungal activity, especially towards to *S. brasiliensis* when compared with *S. globosa*.

A previous multicenter study (27) proposed tentative *S. brasiliensis* ECVs for itraconazole, terbinafine and amphotericin B of 2 µg/mL, 0.12 µg/mL, and 4 µg/mL, respectively. The established ECVs defined here were one dilution higher for itraconazole (4 µg/mL) and amphotericin B (8 µg/mL), whereas for terbinafine the ECV remained unchanged. Additionally, the tentative *S. schenckii* ECVs of itraconazole (4 µg/mL) and amphotericin B (4 µg/mL) were identical to the ECVs in the present evaluation. For *S. globosa* no tentative ECVs have previously been proposed. The established ECV values for itraconazole (4 µg/mL) and amphotericin B (4 to 8 µg/mL) for the three pathogenic *Sporothrix* species are relatively high compared to most clinically significant molds, such as *A. fumigatus* (32). Some however, including *Fusarium* and *Scedosporium* spp. and the Mucorales, are known to have relatively high MICs for these agents.

Importantly, ECVs are intended to distinguish NWT isolates that may harbor resistance mechanisms from wild-type populations of a species. They are solely derived from the natural distribution of MICs and do not incorporate pharmacokinetic/pharmacodynamic parameters or clinical outcome data. As such, they do not predict the therapeutic response; an MIC below the ECV does not necessarily indicate susceptibility to that antifungal agent and clinical success, while an MIC above the ECV does not automatically imply resistance and expected treatment failure. With regard to sporotrichosis, high MICs against itraconazole, based on the previously suggested tentative ECV (2 µg/mL), were not found to correlate with antifungal treatment failure or clinical outcome in humans and cats (23, 26, 31). In cats, none of the 47 feline isolates from Rio Janeiro exhibited high MICs and no association with clinical outcome was observed (23), whereas at the Brazil–Argentina border, therapeutic failure occurred despite low initial MICs (31). This could be a reflection of the pharmacokinetics of itraconazole, which achieves high concentration in the skin, the usual site of infection. Nonetheless, in other reports on *S. brasiliensis* isolates from cats and humans, high MICs (4 to >16 µg/mL) were reported in refractory or more severe cases (24, 28, 33–35).

Using these new defined ECVs, we found that the percentage of NWT isolates varied by fungal species and antifungal drug. The percentage of NWT isolates was low (< 5%) for amphotericin B and terbinafine for all three pathogenic *Sporothrix* species, and was also low for itraconazole and posaconazole for *S. brasiliensis*. Higher percentages of NWT isolates (>5%) were found for *S. schenckii* with 8.3% and 19.5% for itraconazole and posaconazole, respectively, and for *S. globosa* with 7.5% NWT isolates according to tentative ECV for posaconazole. Another study also reported higher itraconazole and posaconazole MICs for *S. schenckii* than for *S. brasiliensis* (36). The NWT isolates, especially those found among *S. schenckii* and *S. globosa* may harbor resistance mechanisms as observed for *A. fumigatus* (37) and *Trichophyton indotineae* (38). However, genes linked to resistance mechanisms for *Sporothrix* species are poorly investigated. Resistance has been correlated with substitutions of the *CYP51* and/or *TAC1* genes in a few *S. brasiliensis* isolates (29, 39), but there are no resistance mechanisms reported that are correlated with the high MIC values of *S. schenckii* or *S. globosa*.

No ECV could be established for isavuconazole or voriconazole, drugs that share a similar structure and activity profile, as MICs were truncated at the high end of the recommended testing range, which suggests potential intrinsic resistance or reduced susceptibility. These results are in agreement with the guideline recommendations against the use of isavuconazole and voriconazole for sporotrichosis treatment (14, 15), which are based on the lack of efficacy of these drugs *in vitro* (34, 40) and in mice (41).

Moreover, no ECVs could be established for olorofim, as data from only one laboratory were available. Olorofim was tested against *S. brasiliensis* and *S. globosa* isolates and showed good *in vitro* activity. In general, *S. brasiliensis* had a lower MIC range (0.004 - 0.25 µg/mL) and MIC_90_ (0.12 µg/mL) against olorofim than *S. globosa* isolates (0.25 - 1 µg/mL; MIC_90_= 1 µg/L). Other studies evaluating olorofim in *A. fumigatus* using the EUCAST methodology found MIC_90_ values (0.125 µg/mL) (42) similar to those of *S. brasiliensis*. Olorofim is the first molecule in a new class of antifungals (orotomides) that inhibit the mitochondrial dihydroorotate dehydrogenase (DHODH) (43). This inhibition impairs pyrimidine biosynthesis leading to decreased cell wall biogenesis; protein glycosylation; the synthesis of DNA, protein, and phospholipids; and consequently, fungal cell death. While this drug is known as a mold-specific agent with little to no activity against yeasts, including *Candida* and *Cryptococcus* spp. among others (44), a previous *in vitro* study showed that olorofim inhibited and killed the yeast forms of *S. brasiliensis*, *S. schenckii*, and *S. globosa* at concentrations lower than itraconazole, and showed anti-biofilm activity (45). Olorofim recently completed a phase II clinical trial for salvage treatment of patients with aspergillosis, rare mold infections and coccidioidomycosis (46), and is now in a phase III clinical trial for invasive aspergillosis (47). Additional laboratory testing and AFST data are needed to calculate species specific *Sporothrix* ECVs for olorofim.

*Sporothrix* spp. are thermally dimorphic fungi and, depending on incubation temperatures of 35°C or 30°C, grow in either the yeast or the mycelial phase respectively. This is important in the context of AFST protocols because conidia are used for inoculation, but the temperature of incubation is conductive to yeast growth. In this study, the CLSI M38 methodology for the mycelial phase was used. Some studies have demonstrated that MIC values of the same isolate were ≥8-fold lower in the yeast phase when compared to the mold phase (24, 27, 31, 34, 48). To obtain a *Sporothrix* culture with a pure yeast phase, a long incubation time (> 2 weeks) at 35°C is needed and confirmation by microscopy is essential (31). A pure mycelial phase is easier to obtain by culturing at 30°C for one week. There is no consensus among experts whether the establishment of ECVs should be based on the mold or yeast phase. While some researchers suggest that the ECVs should be based on the yeast form as this phase is the pathogenic infecting form in the host and might therefore better reflect the clinical outcome (34), others highlight the impractical time to results needed for conversion to the yeast form, which can take up to 2 weeks. Mold phase testing is easier to standardize among different laboratories, which is fundamental for establishing ECV. We also found that the MIC data were more similar among the different participant laboratories with mycelial phase as compared to yeast phase (data not shown).

There is overall agreement that the current *Sporothrix* CLSI mold phase protocol CLSI M38 (32), needs some modifications, since testing at 35°C may lead to a mix of yeast and mold phases. In this study, we found a high level of interlaboratory variation for some antifungals, with a modal MIC not within one or two dilutions to the central modal MIC, leading to MIC data rejection of different laboratories, especially for *S. brasiliensis* with 17 to 30% of laboratories excluded for amphotericin B and azoles and for *S. schenckii* and amphotericin B (31%). This inter-lab variation could be partially explained by incubation at 35°C, inducing a mix of mold and yeast phases, with the yeast phase decreasing the MIC for many antifungals, including itraconazole (34). A solution for this issue would be to revise the CLSI protocol to adjust the incubation temperature to 30°C to ensure that there are no mixed forms for a potentially more standardized, less variable MIC assay. In that case, the ECVs presented here would need to be reevaluated. In addition, it would be imperative to distribute *Sporothrix* spp. reference quality control (QC) isolates with defined MICs to allow laboratories comparative assessment of the standardized method. Unfortunately, there are currently no *Sporothrix* QC strains listed in M38M51S. Further efforts should be made to establish *Sporothrix* QC strains with defined MIC ranges for the three *Sporothrix* species, which should be made available to allow thorough verification and validation of the CLSI broth microdilution methodology for *Sporothrix* species.

A limitation of this study is the inability to establish ECVs for all antifungal drugs evaluated here across the three main pathogenic species of *Sporothrix*. For *S. globosa* ECVs could not be established for all drugs, due to high interlaboratory variation of MICs and many high MIC values. Another limitation is the lack of knowledge regarding the potential presence of both yeast and mold phases, following the CLSI M38 protocol, and whether that caused the high interlaboratory variations. Future studies at different mycology laboratories should test and validate changes in the CLSI M38 protocol to test *Sporothrix* in the pure mycelial phase and establish well characterized *Sporothrix* QC strains to overcome current challenges. MICs of more isolates are needed to confirm the tentative ECVs found here for terbinafine against *S. schenckii* and *S.globosa*.

In summary, following the CLSI M57 methodology for the mycelial phase, we have established official ECVs for *S. brasiliensis* to itraconazole, posaconazole, terbinafine, and amphotericin B; for *S. schenckii* to itraconazole, posaconazole and amphotericin B and a single ECV for *S. globosa* against amphotericin B. These ECVs will facilitate the tracking of NWT strains in the current epidemic in Brazil, and emerging clusters in Southeast Asia. The establishment of an ECV is also the first step towards the systematic collection of laboratory data and possible correlations with clinical data to set antifungal breakpoints.

## Materials and Methods

### Experimental design

Laboratories known to test antifungal susceptibility using the CLSI methodologies were contacted to request MIC data for clinically relevant *Sporothrix* species *(S. brasiliensis, S. globosa* and *S. schenckii)*. Inclusion criteria were that species identification performed using molecular techniques and AFST using the broth microdilution method as outlined in the CLSI reference standard M38 Ed3 for filamentous fungi. MIC data for *S. brasiliensis*, *S. globosa*, and *S. schenckii* isolates were obtained from 19 laboratories on five continents and included in the analysis. In Asia, the laboratories submitting data included: Mycology Division, Department of Medical Microbiology, Post Graduate Institute of Medical Education & Research, Chandigarh, India; National Reference Laboratory for Antimicrobial Resistance in Fungal Pathogens, Vallabhbhai Patel Chest Institute, University of Delhi, Delhi, India; and Peking University First Hospital, Beijing, China. In Europe laboratories included: the National Mycology Reference Laboratory, UK; and the Radboudumc-CWZ Center of Expertise for Mycology, Canisius-Wilhelmina Hospital (CWZ), Nijmegen, The Netherlands. In North America, laboratories submitting data included: the Universidad Autónoma de Nuevo León, Mexico; Johns Hopkins University, USA; the Mycotic Diseases Branch, Centers for Disease Control and Prevention, Atlanta, Georgia, USA; and the Laboratoire de santé publique du Québec, Institut national de santé publique du Québec (INSPQ), Sainte-Anne-de-Bellevue, Québec, Canada. In Oceania, the National Mycology Reference Centre, SA Pathology, Adelaide, Australia submitted data. In South America laboratory submitting data were: the Universidade Federal do Paraná (UFPR), Brazil; the Laboratório de Pesquisa Clínica em Dermatozoonoses em Animais Domésticos (Lapclin-Dermzoo), Instituto Nacional de Infectologia Evandro Chagas (INI), Fiocruz, Brazil; the Universidade Federal de São Paulo (UNIFESP), Brazil; the Universidade Federal do Espírito Santo (UFES), Brazil; the Parasitology and Mycology Center, Instituto Adolfo Lutz, São Paulo, Brazil; the Laboratório de Biologia Celular de Fungos (LBCF) at Universidade Federal do Rio de Janeiro (UFRJ), Brazil; the Universidade Federal do Ceará (UFC), Brazil; the Universidad Nacional del Litoral, Santa Fe city, Argentina; and the Mycology Department of INEIA ANLIS “Dr. C. G. Malbran”, Buenos Aires, Argentina.

### Antifungal susceptibility testing

AFST was performed by broth microdilution as outlined in the CLSI reference standard M38 Ed3 for filamentous fungi (32). The isolates used as reference or quality control strains were *Aspergillus fumigatus* ATCC MYA 3626, *Candida parapsilosis* ATCC 22019 (CBS 604), *Candida krusei (Pichia kudriavzevii)* ATCC 6258 (CBS 573), and *Hamigera insecticola* ATCC MYA-3630; the quality controls were within M38M51S CLSI ranges. Amphotericin B isavuconazole, itraconazole, olorofim, posaconazole, terbinafine and voriconazole were tested. The MICs were determined visually after 48–72 hours (2-3 days, depending on the growth rate of the individual isolate) of incubation at 35°C.The inoculum was adjusted to an absorbance 530 nm at 0.09-0.13 and verified by hematocytometer counting to 0.2 - 2.5 X 10^6^ conidia/mL or with a Cellometer X2 (Nexcelom, Manchester, United Kingdom) to perform cell counts and prepare the inoculum. The antifungal susceptibility endpoints for itraconazole, voriconazole, posaconazole, isavuconazole, olorofim, terbinafine, and amphotericin B were 100% inhibition of growth compared to drug-free control.

### Epidemiological cutoff values

The ECV was established using the iterative statistical method with ECOFFinder (V2.1) with a 97.5% threshold following CLSI M57 ECV guidelines (32, 49). The percent NWT, modal MIC, MIC_90_, and geometric mean were calculated for each antifungal. Tentative ECVs were calculated when the number of isolates was less than 100 and greater than 35. MIC datasets were rejected if they did not conform to CLSI M57 guideline (multimodal, truncated or not within 2 dilutions of the central modal MIC of most laboratories).

### Data disclosure

These ECVs were submitted to vote and were approved by the CLSI subcommittee on antifungal susceptibility tests on January 25th, 2025 and will be published in the next edition of the M57S ECV supplement.

## Acknowledgments

The authors acknowledge the contribution of the members of the ISHAM One-Health Sporotrichosis Working Group for having fostered the communication and collaboration for this study and their leadership role in the prevention and control of zoonotic sporotrichosis. The findings and conclusions in this paper are those of the authors and do not necessarily represent the official position of the U.S. Centers for Disease Control and Prevention.

